# A sensitive period in the neural phenotype of language in blind individuals

**DOI:** 10.1101/592345

**Authors:** Rashi Pant, Shipra Kanjlia, Marina Bedny

## Abstract

In congenital blindness, “visual” cortices respond to linguistic information, and fronto-temporal language networks are less left-lateralized. Does this plasticity follow a sensitive period? We tested this by comparing the neural basis of sentence processing in two experiments with adult-onset blind (AB, *n*=16), congenitally blind (CB, *n*=22) and blindfolded sighted controls (*n*=18). In Experiment 1, participants made semantic judgments for spoken sentences and solved math equations in a control condition. In Experiment 2, participants answered “who did what to whom” questions for grammatically complex (with syntactic movement) and grammatically simpler sentences. In a control condition, participants performed a memory task with lists of non-words. In both experiments, visual cortices of CB and AB but not sighted participants responded more to sentences than control conditions, but the effect was much larger in the CB group. Crucially, only the “visual” cortex of CB participants responded to grammatical complexity. Unlike the CB group, the AB group showed no reduction in left-lateralization of fronto-temporal language network relative to the sighted. These results suggest that blindness during development modifies the neural basis of language, and this effect follows a sensitive period.

## INTRODUCTION

When it comes to neural and cognitive development of language, timing is of the essence. Young children acquire language rapidly and effortlessly, without explicit training (Bonvillian, Orlansky, Novack, & Folven, 2012; Gleitman & Wanner, 1982; Petitto, Holowka, Sergio, & Ostry, 2001; Petitto, Katerelos, et al., 2001; Gleitman & Newport, 1995; Lightbrown & Spada, 1993). By contrast, when a second language is acquired in adulthood, learning proceeds more slowly, and plateaus at lower levels of proficiency in phonology and morphosyntax (Johnson & Newport, 1989, 1991; Neville, Mills, & Lawson, 1992, Newport, Bavelier, & Neville, 2001). Deaf individuals who do not have access to sign language until later in life do not attain the same level of proficiency as native signers (Emmorey, Bellugi, Friederici, & Horn, 1995; Mayberry, Chen, Witcher, & Klein, 2011; Mayberry & Eichen, 1991; Mayberry & Lock, 2003).

There is also evidence that the neural systems that support language are particularly plastic early in life (MacSweeney, Capek, Campbell, & Woll, 2008; MacSweeney, Waters, Brammer, Woll, & Goswami, 2008; Mayberry et al., 2011). Delays in language acquisition modify the neural basis of language processing (Neville et al., 1998; Allen, Emmorey, Bruss, & Damasio, 2013; MacSweeney, Waters, et al., 2008; Mayberry & Kluender, 2018). Furthermore, unlike adults, children suffering from early damage to left hemisphere language networks have language processing abilities in the normal range, and recruit right-hemisphere homologues of left-hemisphere fronto-temporal language regions during language tasks (Dronkers, Wilkins, Van Valin, Redfern, & Jaeger, 2004, Rasmussen & Milner, 1977; Zevin, Datta, & Skipper, 2012; Newport et al., 2017; Kempler, Van Lancker, Marchman, & Bates, 1999; Rosen et al., 2000; Tivarus, Starling, Newport, & Langfitt, 2012).

Consistent with this prior evidence for the malleability of the neural basis of language during development, evidence from studies of blindness suggests that the language network can be augmented with cortical real-estate that is typically occupied by visual perception. Individuals who are blind from birth recruit a network of “visual” areas during sentence processing, lexical retrieval, reading and word production tasks (Hamilton & Pascual-Leone, 1998; Kupers et al., 2007; Sadato et al., 1998; Bedny, Pascual-Leone, Dodell-Feder, Fedorenko, & Saxe, 2011; Lane, Kanjlia, Omaki, & Bedny, 2015; Lane et al., 2017; Röder, Stock, Bien, Neville, & Rösler, 2002; Watkins et al., 2012). This recruitment is part of a broader phenomenon, whereby in blindness, regions of the “visual” cortex are recruited for non-visual functions, including spatial localization and numerical cognition (for review, see Bedny, 2017; Collignon et al., 2011; Kanjlia, Lane, Feigenson, & Bedny, 2016; Kim, Kanjlia, Merabet, & Bedny, 2017; Röder, Rösler, & Neville, 2000). Language-responsive “visual” cortices are selectively involved in language-processing, rather than all auditory and tactile tasks (e.g. respond more to sentences than math equations) (Kanjlia, Lane, Feigenson, & Bedny, 2016). These “visual” language regions are, furthermore, sensitive to high-level linguistic information i.e. semantics and grammar. They respond more to sentences than Jabberwocky, and more to Jabberwocky than lists of unconnected non-words (Bedny et al., 2011; Lane et al., 2015; Röder et al., 2000). Responses are also higher for grammatically complex than simpler sentences (Röder et al., 2002, Lane, et al., 2015). Finally, language-responsive “visual” areas are co-lateralized with the fronto-temporal language network, and show higher functional correlations with classical “language” regions even in the absence of a task (i.e. at rest) (Kanjlia, Lane, Feigenson, & Bedny, 2016; Bedny, et al., 2011; Lane et al., 2017; Liu et al., 2007).

One hypothesis is that in blindness, parts of “visual” cortex are incorporated into the language system during development, when cortical specialization is taking place (Bedny, Richardson, & Saxe, 2015). According to the developmental specialization hypothesis, absence of visual input during a sensitive period enables “visual” cortices to develop specialization for language processing. Alternatively, it remains possible that “visual” cortex has a latent ability to respond to linguistic information in all humans, irrespective of developmental visual history. These accounts make different predictions with respect to blindness onset. The developmental specialization hypothesis predicts that “visual” cortex responses to language are particular to congenital blindness. By contrast, the unmasking hypothesis predicts that “visual” cortex recruitment for language should also occur in people who lose their vision as adults.

The available evidence is mixed with regard to whether language-related visual cortex plasticity follows a sensitive period. Some activity has been observed during language tasks in late blind adults, and even in sighted blindfolded participants (Büchel, Price, Frackowiak, & Friston, 1998; Burton, Diamond, & McDermott, 2006; Burton, Snyder, Diamond, & Raichle, 2006; Burton et al., 2002; Elli, Lane, & Bedny, 2019). For example, like congenitally blind individuals, late-onset blind individuals activate “visual” cortices during Braille reading and verb generation (Büchel et al., 1998; Burton et al., 2002; Burton, Diamond, et al., 2006; Burton & McLaren, 2006). One study of resting state connectivity found that individuals with retinitis pigmentosa, who did not become totally blind until adulthood, show elevated correlations between inferior frontal language areas and occipital cortices (Sabbah et al., 2016). However, such resting state changes are much larger in individuals blind from birth (Kanjlia, Pant, & Bedny, 2018).

On the other hand, there is also some evidence that “visual” cortex activity during language tasks in congenitally and late-onset blind individuals may be different. Some studies find that late and congenitally blind people activate different parts of the “visual” cortex during Braille reading (Büchel et al., 1998; Burton et al., 2006; Burton et al., 2002). One study compared activity during sentence processing and a working memory task with meaningless sounds in congenitally blind and late-onset blind individuals. This study found higher responses during the sentence processing task only in people born blind (Bedny, Pascual-Leone, Dravida, & Saxe, 2012).

A key outstanding question is whether “visual” cortices of late blind individuals, like those of people who are born blind, show signature responses to higher-order linguistic information, such as syntax and, if so, whether they do so to the same degree. All prior studies with late-onset blind individuals have compared language tasks to relatively low-level control conditions. It therefore remains possible that activation during language tasks in congenitally blind and late-onset blind individuals reflects different cognitive operations.

The goal of the present study was to ask whether blindness-related recruitment of the “visual” cortex for language processing, and development of sensitivity to grammatical structure in particular, follows a sensitive period of development. To this end, we compared the neural basis of language in adult-onset blind, congenitally blind and sighted individuals in two experiments. Experiment 1 compared spoken sentence comprehension to an auditory math task. Experiment 2 compared sentences to lists of non-words, and manipulated the grammatical complexity of the sentences using a syntactic movement dependency, while holding lexical semantics constant. As noted above, a previous study showed that regions in the “visual” cortex of congenitally blind individuals responds more to sentence-processing than control tasks, and more to grammatically complex than grammatically simple sentences. The goal of the current study was to ask whether visual cortices of adult-onset blind individuals show similar signatures of linguistic sensitivity.

The present study also allowed us to test a second question. In addition to the recruitment of “visual” areas for language, congenital blindness is also associated with reduced left-lateralization of front-temporal language areas themselves (Lane et al., 2015; Lane et al., 2017; Röder et al., 2002). This phenomenon appears to be unrelated to the recruitment of visual cortex for language per se, since across blind individuals the amount of “visual” cortex recruitment for language is not predictive of the laterality of fronto-temporal language networks. Furthermore, although there is some evidence that recruitment of the “visual” cortex for language processing confers cognitive benefits, reduction in left lateralization appears to have no consequences for behavior (Lane et al., 2015; Lane et al., 2017). The goal of the current study was to ask whether language laterality is also changed in adult-onset blind individuals. Based on previous work showing sensitive period effects in language acquisition, we hypothesized that modification of the language network in blindness, including reduced left-lateralization and recruitment of the visual cortex, follows a sensitive period.

One challenge in answering the question of whether there is a sensitive period for the effects of vision loss is determining the relevant cut off point for “late” blindness. Previous studies have defined late blindness in various ways, including vision loss starting at 7, 9, 11 and 16 years of age. In the current study, we took a conservative approach - the “late” blind group includes only participants who lost their vision at 17 years of age or later. Therefore, we henceforth refer to this group as “adult-onset blind”. The congenitally blind group includes participants who had at most minimal light perception from birth. If sensitive period effects are observed in the current study, in future work it will be important to test participants who lost their vision at various ages during childhood, to empirically define the sensitive period of development.

## METHODS

### Participants

Sixteen adult-onset blind individuals (AB; 5 female, mean age = 56.87, SD age = 10.39, mean years of education = 17.31, SD years of education = 3.11), twenty-two congenitally blind (CB; 16 female, mean age = 46.50, SD age = 17.18, mean years of education = 16.67, SD years of education = 2.26) and eighteen blindfolded sighted controls (S; 9 female, mean age = 46.50, SD age = 15.32, mean years of education = 16.34, SD years of education = 1.37) contributed data to the current study. Adult-onset blind participants were blind for at least 4 years, with a mean blindness duration of 16 years. One adult-onset blind participant only contributed data to Experiment 1. This participant did not learn English until 11 years of age, and was therefore excluded from data analyses of Experiment 2, which manipulated syntactic complexity. One additional adult-onset blind participant acquired English at 5 years-of-age, however, as their acquisition was early and their performance was not different from the group, they were included in both experiments. We additionally excluded any scanned participant who performed below 55% on the sentence condition of either experiment (chance = 50%). This resulted in exclusion of 3 congenitally blind participants, not included in the subject count.

For both groups of blind individuals, all causes of blindness were related to pathology of the retina or optic nerve, not brain damage (Table 1). Adult-onset blind participants were fully sighted until 17 years of age or later (vision loss between the ages of 17 to 70, mean = 33.19, SD = 12.81, Table 1). At the time of the experiment, all blind participants had at most minimal light perception (LP) or no light perception (NLP) and the proportion of participants with light perception did not differ across blind groups (proportion with light perception AB 38%, CB 45%). Braille reading ability of participants was self-reported via a score on a scale of 1-5 (AB average = 2.5, CB average = 4.82). The question was worded as: On a scale of 1 to 5, how well are you able to read Braille, where 1 is “not at all”, 2 is “very little”, 3 is “reasonably well”, 4 is “proficiently”, and 5 is “expert”. None of the participants suffered from any known cognitive or neurological disabilities. All participants gave written informed consent and were compensated $30 per hour. (Data from 19 CB and 18 S participants have previously been described in Lane et al., 2015.)

**Table 1:** Participant demographic information and vision loss history. For adult-onset blindness, duration of blindness was calculated by subtracting age at which current level of vision was reached from age at time scanned (Mean = 16.06 years SD = 10.58 years). For congenital blindness, duration of blindness is the age when scanned (Mean = 46.50 years SD = 17.18). Most common causes of adult-onset blindness were Diabetic Retinopathy (DR), Glaucoma and Macular Degeneration (GMD) and Retinitis Pigmentosa (RP). Most common causes of congenital blindness were Leber’s Congenital Amaurosis (LCA) and Retinopathy of Prematurity (ROP), optic nerve neuropathy (ONN).

| Subject | Cause of Blindness | Current Level of Vision | Age at time of scan | Age of Blindness Onset | Age reached current level of vision | Blindness Duration | Braille Reading Score |
| --- | --- | --- | --- | --- | --- | --- | --- |
| LB1 | Autoimmune | NLP | 61 | 37 | 57 | 4 | 2 |
| LB2 | Trauma | LP | 45 | 22 | 22 | 23 | 1 |
| LB3 | DR | NLP | 48 | 17 | 17 | 31 | 3 |
| LB4 | RP | LP | 53 | 33 | 35 | 18 | 3 |
| LB5 | Trauma | LP | 49 | 19 | 19 | 30 | 4 |
| LB6 | RP | NLP | 34 | 17 | 25 | 9 | 3 |
| LB7 | GMD | LP | 67 | 48 | 49 | 18 | 2 |
| LB8 | DR | NLP | 66 | 45 | 47 | 19 | 2 |
| LB9 | RP | LP | 64 | 28 | 59 | 5 | 1 |
| LB10 | GMD | NLP | 69 | 49 | 59 | 10 | 3 |
| LB11 | RP | LP | 74 | 38 | 70 | 4 | 1 |
| LB12 | ONN | NLP | 50 | 21 | 34 | 16 | 2 |
| LB13 | GMD | NLP | 51 | 38 | 38 | 13 | 1 |
| LB14 | Trauma | NLP | 59 | 50 | 50 | 9 | 5 |
| LB15 | DR | NLP | 60 | 19 | 20 | 40 | 5 |
| LB16 | DR | NLP | 60 | 50 | 52 | 8 | 2 |
| CB1 | LCA | LP | 22 | -- | -- | 22 | 5 |
| CB2 | ROP | LP | 32 | -- | -- | 32 | 4 |
| CB3 | ROP | LP | 70 | -- | -- | 70 | 4 |
| CB4 | Unknown | NLP | 43 | -- | -- | 43 | 5 |
| CB5 | ROP | LP | 67 | -- | -- | 67 | 5 |
| CB6 | ROP | NLP | 67 | -- | -- | 67 | 5 |
| CB7 | ROP | LP | 26 | -- | -- | 26 | 4 |
| CB8 | ROP | NLP | 64 | -- | -- | 64 | 5 |
| CB9 | LCA | LP | 35 | -- | -- | 35 | 5 |
| CB10 | LCA | NLP | 47 | -- | -- | 47 | 4 |
| CB11 | ROP | NLP | 39 | -- | -- | 39 | 5 |
| CB12 | LCA | LP | 49 | -- | -- | 49 | 5 |
| CB13 | LCA | LP | 25 | -- | -- | 25 | 5 |
| CB14 | ROP | NLP | 62 | -- | -- | 62 | 5 |
| CB15 | ROP | NLP | 62 | -- | -- | 62 | 5 |
| CB16 | ROP | NLP | 60 | -- | -- | 60 | 5 |
| CB17 | ROP | NLP | 46 | -- | -- | 46 | 5 |
| CB18 | ROP | NLP | 61 | -- | -- | 61 | 5 |
| CB19 | ROP | NLP | 67 | -- | -- | 67 | 5 |
| CB20 | LCA | LP | 28 | -- | -- | 28 | 5 |
| CB21 | LCA | LP | 19 | -- | -- | 19 | 5 |
| CB22 | ONN | NLP | 32 | -- | -- | 32 | 5 |

### Experimental Procedures

Participants were scanned while performing two separate auditory tasks. All stimuli were presented over Sensimetrics MRI compatible earphones (http://www.sens.com/products/model-s14/). Volume was adjusted to a comfortable level for each participant. All participants, blind and sighted, wore a blindfold for the duration of the experiment.

#### Experiment 1 (Sentences and Equations)

Experiment 1 consisted of a language task and a mathematical control task (Kanjlia et al., 2016, Lane et al., 2015). In the language task, participants judged whether the meanings of two consecutively presented sentences, one presented in active voice and one in passive voice, were the same. For the “same” trials, the relations and roles of the people in the sentences were maintained. On the “different” trials, the roles were reversed (e.g. “The bartender that the mailman knew cut the grass” and “The grass was cut by the mailman that the bartender knew.”).

In the mathematical control task, participants judged whether the value of ‘X’ in two consecutively presented subtraction equations was the same. ‘X’ could occur as either the operand (e.g.: 6 – X = 3) or answer (e.g.: 16 – 13 = X). The equations varied in difficulty level, however, the difficulty manipulation was not analyzed in the present experiment (see Kanjlia et al., 2016 for further details).

There were 48 sentence trials and 96 mathematical trials. Each trial was 14 s long, starting with a 0.25 s tone followed by two sentences/equations of 3.5 s each, separated by a 2.75 s interval. After hearing the second stimulus, participants had 4 s to respond. The experiment included 36 rest blocks that were 16 s long.

#### Experiment 2 (Sentences and Nonwords)

In Experiment 2, participants performed a sentence processing task, and a non-word working memory control task. In the sentence task, participants listened to a sentence, followed by a yes or no question, which required participants to judge who did what to whom. Half of the sentences were more syntactically complex (MOVE) and half were less complex (NONMOVE). The MOVE sentences contained a syntactic movement dependency in the form of an object-extracted relative clause (e.g.: “The accountant [that the corrupt detective in the organized crime division dislikes] advises the Sicilian mob.”). Sentences with movement require listeners to relate distant elements (words and phrases) to each other during the derivation of the sentence’s structure (Chomsky, 1957). The NONMOVE sentences had similar meanings and contained nearly identical words to the MOVE sentences, but did not contain an object extracted relative clause (e.g.: ‘The corrupt detective in the organized crime division dislikes [that the accountant advises the Sicilian mob.]’). Sentences were yoked across conditions, such that each sentence had both a MOVE and a NONMOVE version. Each participant heard one version of the sentence, counterbalanced across participants.

In the non-word working memory control task, participants heard a long list of non-words (target), followed by a shorter list of non-words (probe) which consisted of non-words from the first list - either in the same order as they were initially presented, or in a different order. Subjects judged whether the non-words in the shorter probe list were in the same order as they had occurred in the initially presented, longer target list.

There were 54 trials each of the MOVE, NONMOVE and NONWORD conditions divided across 6 runs, i.e. 9 in each run. All the trials were 16 s long, consisting of a tone, a 6.7 s sentence/target non-word list, 2.9 s question/probe non-word list, giving participants until the end of the 16s periods to respond. We matched the sentences and target non-word sequences for number of items (words and nonwords, sentence = 17.9, nonword lists = 17.8; *p =* 0.3), number of syllables per item (sentence = 1.61, nonword = 1.59; *p =* 0.3), and mean bigram frequency per item (sentence = 2.34, nonword = 2.35; *p =* 0.3) (Duyck, Desmet, Verbeke, & Brysbaert, 2004). For further details, see Lane et al., 2015.

### MRI ACQUISITION AND DATA ANALYSIS

MRI structural and functional scans were acquired on a 3 Tesla Phillips MRI. For the structural T1 weighted images, 150 axial slices with 1mm isotropic voxels were collected, and for the functional BOLD images, 36 axial slices with 2.4 × 2.4 × 3 mm voxels were collected with TR 2 seconds.

We created cortical surface models for each subject using the Freesurfer pipeline, and used FSL, Freesurfer, HCP workbench and custom software for surface-based analyses. Functional data were motion corrected, high pass filtered (128 s cutoff), resampled to the cortical surface and smoothed with a 6 mm FWHM Gaussian kernel on the cortical surface. Only cortical data, excluding the cerebellum and subcortical structures, was analyzed. BOLD activity as a function of condition was analyzed using a GLM and combined across runs within subjects using fixed-effects analyses. For both experiments, predictors were entered after convolving with a canonical HRF and its first temporal derivative. In Experiment 1, each type of math and language trial was a separate predictor. For Experiment 2, the non-word trials and each kind of sentence trial were separate predictors. We dropped trials where the participants failed to respond by including a regressor of no interest (average drops per run CB=1.21, AB=1.32, S=1.38). We also dropped time-points with excessive (>1.5 mm) motion. Data was combined across participants using random effects analysis. We used cluster correction in all whole brain analyses in order to correct for multiple comparisons across the cortical surface at p < 0.05.

### ROI ANALYSES

Region of interest (ROI) analyses were used to probe responses to language in the visual cortices of adult-onset blind participants and compare them to those of the congenitally blind and blindfolded sighted participants. A two-step procedure was used to define individual-subject specific functional ROIs (Fedorenko, Hsieh, Nieto-Castañón, Whitfield-Gabrieli, & Kanwisher, 2010; Saxe, Brett, & Kanwisher, 2006). First, visual cortex search-spaces were defined based on a combination of anatomical landmarks, previous literature and orthogonal group-wise contrasts. Next, individual subject orthogonal ROIs were defined within these search-spaces by either using data from one experiment to select task-responsive vertices and extracting data from the other, or performing a leave-one-run out procedure, described in detail below.

To examine responses to sentences relative to math equations (Experiment 1), we first used a leave-one-run-out procedure within an anatomically defined V1 search-space. The V1 search-space was defined in each individual participant based on sulcal and gyral landmarks, according to previously published procedures (Hadjikhani, Liu, Dale, Cavanagh, & Tootell, 1998; Van Essen, 2005). Within this search space, we selected the top 20% most responsive vertices to sentences > equations for each subject. Vertices were selected based on data from all but one run, and PSC was extracted from the left-out run, iteratively over all possible leave-one-out combinations. The results of each leave-one-out procedure were averaged together.

Second, we looked for a sentences > equations effect (Experiment 1) in those visual cortex regions that responded more to sentences than nonwords in the adult-onset blind group as compared to the sighted in Experiment 2. A group-wise searchspace was defined as adult-onset blind > sighted for sentences > nonwords, at a leninent threshold of p < 0.01, uncorrected (AB language responsive visual cortex region, AB LangOccip). This search-space was then truncated anteriorly using the PALS atlas occipital lobe boundary (Van Essen, 2005). Within this search-space, we performed the same leave-one-run out procedure as described above to define individual-subject functional ROIs using the sentences > equations contrast from Experiment 1.

To test for the sentences > nonwords and grammatical complexity effects in Experiment 2 we used three ROIs. First, we examined activity in V1. An anatomical V1 search-space was defined as described above. Within this search-space, orthogonal individual subject functional ROIs were defined for each participant using the sentence > equations contrast from Experiment 1 (top 20% sentences > equations). No leave-one-run-out procedure was necessary, since Experiment 1 data were used to define ROIs for Experiment 2 analyses.

Second, we examined activity in visual areas that have previously been found to respond to spoken language in those who are congenitally blind (Kanjlia et al., 2016; Kim et al., 2017; Lane et al., 2015). A search space was created using the group-wise data from Experiment 1, defined as the occipital cortex regions that responded to sentences > equations (p < 0.05) in the congenitally blind more than sighted group (CB language responsive occipital cortex region – CB LangOccip). This search-space was then truncated anteriorly using the PALS atlas occipital lobe boundary (Van Essen, 2005). Next, within each search-space, we defined individual-subject-specific functional ROIs by choosing the top 20% of vertices that showed the sentences > equations effect for that particular subject. Again, no leave-one out procedure was necessary because ROIs were defined based on an independent experiment.

To further probe for the grammatical complexity effect, we also conducted an ROI analysis within the AB language responsive occipital cortex region (AB LangOccip), that was more responsive to sentences than nonwords in the adult-onset blind than sighted group. This analysis was conducted to ensure that the grammatical complexity effect was not missed in the adult-onset blind group by focusing on regions that were more relevant to the congenitally blind group. Individual subject functional ROIs were defined within the AB occipital search-space by taking the top 20% of sentences > equations responsive vertices contrast from Experiment 1.

All of the above search-spaces and contrasts used were orthogonal with respect to the contrasts of interest. Note however that the AB language responsive visual cortex search-space is specifically looking at parts of the visual cortex that respond to spoken language in the AB group more so than in the sighted, whereas the CB language responsive visual cortex search-space focuses on areas that are more responsive to language in those who are congenitally blind relative to the sighted. These approaches are therefore complementary to each other, ensuring that no effects are missed because of different visual cortex areas recruited for language in these two populations. In practice, these approaches yield similar results, suggesting that similar visual cortex regions become responsive to language in congenitally and adult-onset blind individuals.

A classically language responsive ROI was defined using a similar procedure to the individual subject visual cortex CB occipital functional ROIs. Within the Inferior Frontal Gyrus (IFG) search space from Fedorenko et al., we selected the top 20% most responsive vertices to Experiment 1 (sentences > equations) in each individual subject, and examined responses to Experiment 2 (Fedorenko et al., 2010). All ROIs were defined in both hemispheres. Previous studies have found reduced left lateralization of language in congenitally blind individuals (Lane et al., 2017; Röder, Rösler, & Neville, 2000). Whether lateralization is also reduced in adult-onset blindness is not known. To account for potential lateralization differences across congenitally blind, adult-onset blind and sighted participants we conducted analyses in every subject’s language dominant hemisphere (see Lane et al., 2015 for similar analysis). For each participant, we calculated (L-R)/(L+R), where L and R are the sum of positive z-statistics > 2.3 (p < 0.01 uncorrected) in the left and right hemisphere, respectively (Lane et al., 2015; Lane et al., 2017). Laterality was defined based on the entire hemisphere, minus the occipital lobe. This was done to avoid biasing laterality indices based on visual cortex plasticity differences across groups. We used data from Experiment 1 to determine the laterality index and then analyzed results from Experiment 2, and vice versa.

For all of the above ROIs, PSC was calculated as BOLD signal during the predicted peak window (8-14 s for Experiment 1, 6-12 s for Experiment 2) relative to rest ((Signal condition - Signal baseline)/Signal baseline). PSC was averaged across vertices within each ROI.

## Results

### 1. Behavioral Results

#### 1.1 Experiment 1

Adult-onset blind participants were as accurate on the language as compared to the math condition (t(15) = 0.26, p = 0.800), and marginally faster on the language trials (t(15) = 2.18, p = 0.052). There were no significant differences between any of the three groups in their accuracy on the math trails (One way ANOVA effect of group F(2,52) = 2.02, p = 0.143) or the sentence trials (One way ANOVA effect of group F(2,52) = 1.3, p = 0.282). The congenitally blind group was faster on the language trials than both the adult-onset blind (t(15) = 2.24, p = 0.036) and the sighted group (t(17) = 2.06, p = 0.046), who were not different from each other (t(15) = − 0.44, p = 0.662). There was no difference in any of the three groups in their RT on math trials (One way ANOVA effect of group F(2,52) = 2.10 p = 0.133)

#### 1.2 Experiment 2

Like congenitally blind and sighted participants, adult onset blind participants were less accurate and slower on the MOVE than NONMOVE sentences (Figure 1, AB: Accuracy t(14) = −6.77, p < 0.001; RT: t(14) = 7.01, p < 0.001 CB: Accuracy t(21) = −5.35, p < 0.001; RT t(21) = 6.06, p < 0.001 S: Accuracy t(17) = − 7.69, p < 0.001 RT t(17) = 5.26, p < 0.001). The effect of movement in the adult-onset blind was no different from either the congenitally blind or the sighted (group-by-condition ANOVA, group-by-condition interaction: Accuracy CB vs AB: F(1,35) = 1.25, p = 0.272; Accuracy AB vs S: F(1,31) = 0.25, p = 0.619; RT CB vs AB: F(1,35) = 0.95, p = 0.335; RT AB vs S: F(1,31) = 1.84, p = 0.185). The movement effect was also not different when comparing the congenitally blind and sighted groups (group-by-condition ANOVA, group-by-condition interaction: Accuracy CB vs S: F(1,38) = 1.92, p = 0.174; RT CB vs S: F(1,38) = 0.16, p = 0.695)

**Figure 1:**
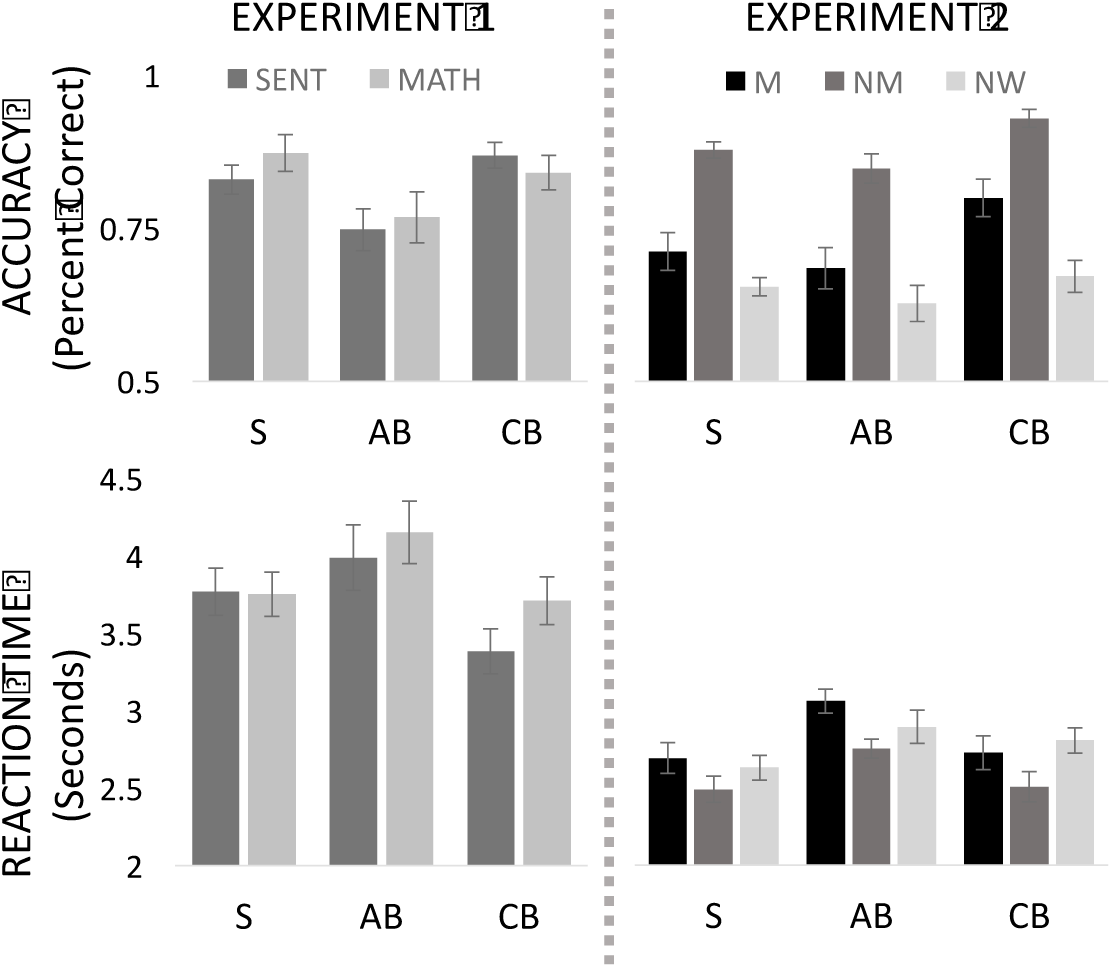
Behavioral performance of Sighted (S), Adult-onset Blind (AB) and Congenitally Blind (CB) on Experiment 1 (Sentence (SENT) and Mathematical equations (MATH)) and Experiment 2 (MOVE (M), NONMOVE (NM) and NONWORD (NW)) conditions. Error bars represent standard error of mean (SEM).

Across sentence types, the adult-onset blind participants were no different in their accuracy from the sighted (group-by-condition ANOVA main effect of group AB vs S F(1,31) = 0.53, p = 0.474) but were significantly less accurate than the congenitally blind (group-by-condition ANOVA main effect of group CB vs AB F(1,35) = 10.77, p = 0.002) Congenitally blind participants were also more accurate than the sighted group (group-by-condition ANOVA main effect of group CB vs S F(1,38) = 6.91, p = 0.012).

In reaction time, the adult-onset blind group was slightly slower at the sentence comprehension task than both the sighted and congenitally blind groups (group-by-condition ANOVA main effect of group AB vs S F(1,31) = 5.18, p = 0.030; CB vs AB F(1,35) = 3.66, p = 0.063), which were not different from each other (group-by-condition ANOVA main effect of group CB vs S F(1,38) = 0.02, p = 0.889).

For the non-word condition, there was no difference between the three groups in accuracy or response time in a one-way ANOVA. (One way ANOVA effect of group: Accuracy F(2,52) = 0.77, p = 0.467; RT F(2,52) = 1.37, p = 0.264).

Additionally, we repeated all behavioral analyses with age matched subsets (S Mean Age = 53.07, SD Age = 9.80, n = 14; AB Mean Age = 54.78, SD Age = 9.27 n= 14; CB Mean Age = 54.43, SD Age = 12.65, n = 16; t-tests between distributions p > 0.1) and the results did not change.

### 2. fMRI Results

#### 2.1 Larger responses to language in “visual” cortex of congenitally than adult-onset blind individuals (Experiment 1)

##### Whole-brain analysis

Congenitally blind, but not sighted participants show larger responses to sentences than mathematical equations in lateral occipital and posterior fusiform cortices (p < 0.05, within CB group and CB > S group-by-condition interaction, cluster corrected, Figure 2). In adult-onset blind participants, occipital responses did not reach significance in this contrast (Figure 2, AB > S group-by-condition interaction).

**Figure 2:**
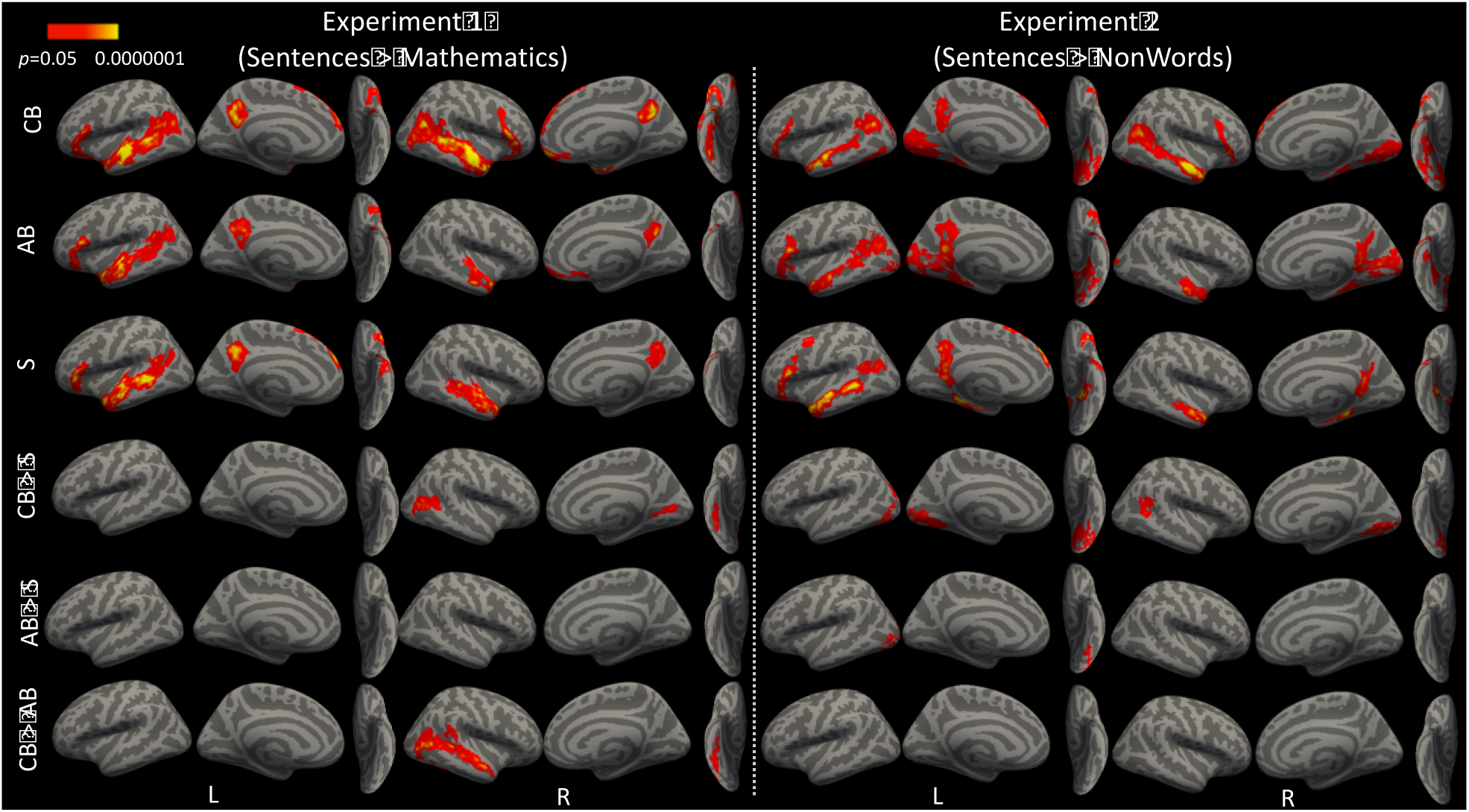
Whole brain analysis results of all subjects in the left and right hemispheres on the lateral, medial and ventral surface (cluster corrected, p < 0.05) for Experiment 1 (left) and Experiment 2 (right) in the Congenitally Blind (CB, n=22), Adult Onset Blind (AB, n=16 (Experiment 1), 15 (Experiment 2)) and Sighted (S, n=18) groups.

##### ROI analysis

Since whole-brain analyses could potentially miss small effects in the adult-onset blind group, we used a sensitive individual-subject leave-one-run out ROI analysis to look for responses to sentences more than equations in the adult-onset blind group.

The response to sentences compared to equations was larger in the congenitally blind than the sighted group in both visual cortex ROIs (group-by-condition ANOVA CB vs S group-by-condition interaction LangOccip: F(1,38) = 12.97, p < 0.001, LangV1: F(1,38) = 8.79, p = 0.005). By contrast, the difference between the adult-onset blind group and the sighted group did not reach significance in either secondary visual areas or in V1 (group-by-condition ANOVA AB vs S group-by-condition interaction LangOccip: F(1,32) = 0.53, p = 0.471; LangV1: F(1,32) = 2.30, p = 0.139).

When adult-onset blind adults were compared to congenitally blind adults directly, the response to sentences (relative to equations) was smaller in the adult-onset blind group in secondary visual areas (LangOccip: group-by-condition ANOVA CB vs AB group-by-condition interaction F(1,36) = 6.94, p = 0.012) but was not different from congenitally blind individuals in V1 (LangV1: group-by-condition ANOVA CB vs AB main effect of condition F(1,36) = 22.61, p < 0.001; group-by-condition interaction F(1,36) = 2.54, p = 0.119).

In post-hoc t-tests, we found a larger response to sentences than equations in the congenitally blind group in both visual cortex ROIs (LangOccip: t(21) = 6.95, p < 0.001; LangV1: t(21) = 4.09, p < 0.001). In the sighted group, a significant effect was present in the LangOccip ROI (t(17) = 2.74, p = 0.014) but there was no significant effect for sentences > equations in V1 (t(17) = 0.85, p = 0.407). In adult-onset blind individuals, the effect was present in in both V1 and secondary visual areas (LangV1: t(15) = 2.043, p = 0.028; LangOccip: t(15) = 3.09, p = 0.007) (Figure 3).

**Figure 3:**
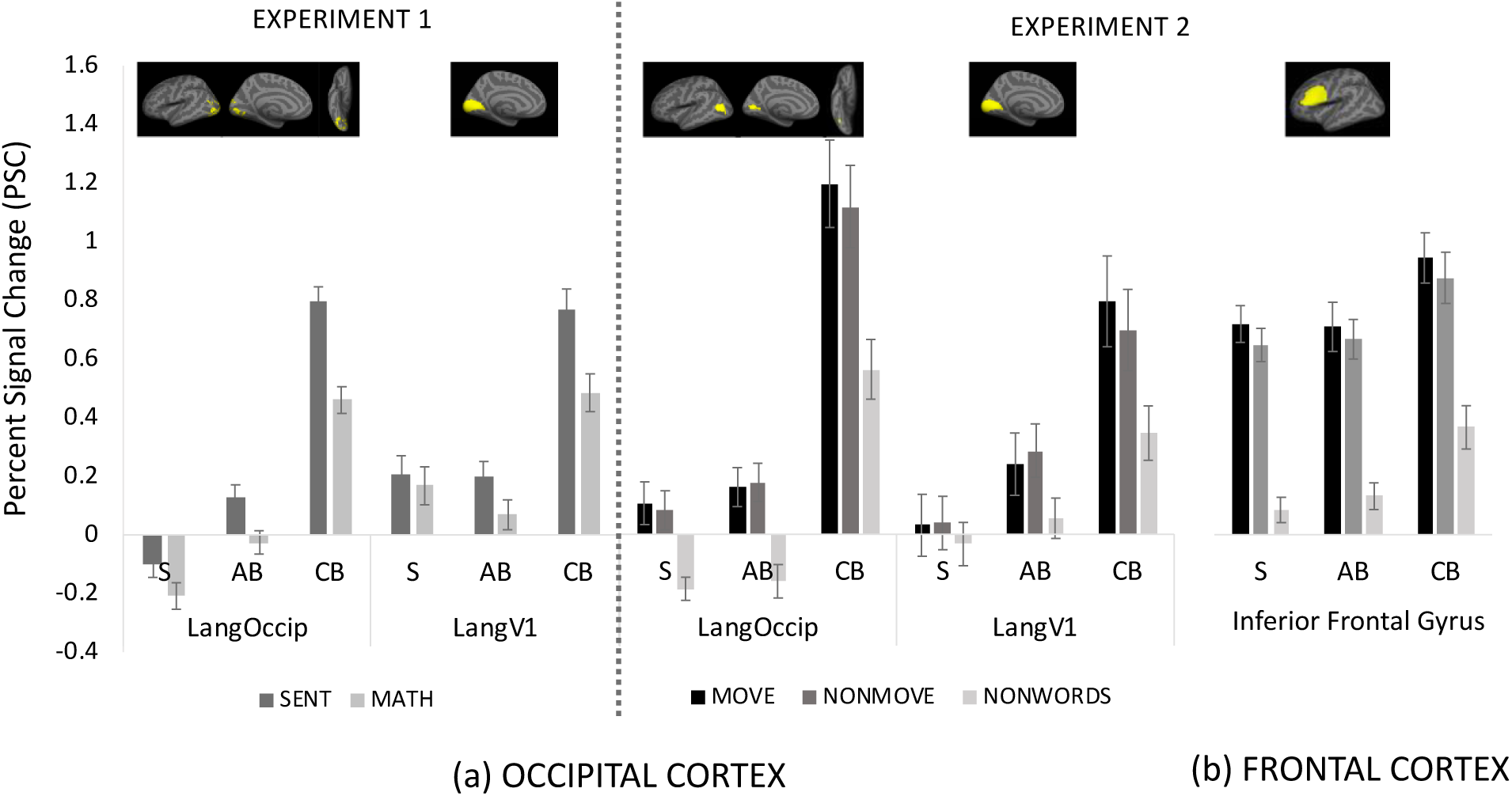
Percent Signal Change (PSC) in response to the Sentence and Equation conditions of Experiment 1 and the MOVE, NONMOVE and NONWORD conditions of Experiment 2 in the Sighted (S), Adult-Onset blind (AB) and Congenitally Blind (CB) groups in (a) occipital cortex and (b) frontal cortex ROIs. Error bars represent SEM.

The overal response relative to rest was also larger in congenitally blind as compared to the adult-onset blind individuals. In the LangOccip ROI, the adult-onset blind individuals fell intermediate between sighted and congenitally blind (LangOccip CB vs AB: F(36) = 28.95, p < 0.001; AB vs S: F(32) = 7.05, p = 0.012). In V1, adult-onset blind participants were no different from the sighted (LangV1 CB vs. AB F(1,36) = 10.10, p = 0.003; AB vs S F(32) = 0.30, p = 0.585).

In sum, the “visual” cortices of adult-onset blind participants showed a smaller response to language (i.e. sentences > math) relative to congenitally blind participants. However, in some “visual” regions, responses to language were observed even in the adult-onset blind group.

### 2.2 “Visual” cortex of adult-onset blind individuals responds to spoken sentences more than to lists of non-words, but less so than in congenitally blind adults (Experiment 2)

#### Whole-brain analysis

There were larger responses to sentences than nonwords in the visual cortex of congenitally blind but not sighted participants in lateral occipital cortex bilaterally, retinotopic visual cortices on the medial surface (in the location of V1, V2 and V3) as well as the posterior fusiform on the ventral surface (CB > S group-by-condition interaction, p < 0.05, cluster corrected) (Figure 2).

The visual cortices of adult-onset blind participants showed a qualitatively similar but weaker response to spoken sentences compared to the congenitally blind group. Larger responses to spoken language than non-words was observed in posterior lateral occipital cortex, within the vicinity of the lateral occipital complex (LO) and V5 (MT/MST). Small patches of activation were also present on the medial surface in pericalcarine and extrastriate cortices (in the regions of V1, V2 and V3). In a group-by-condition interaction analysis, we observed larger visual cortex responses to sentences than nonwords in the adult-onset blind as compared to the sighted in the posterior lateral occipital cortex (AB > S, p < 0.05, cluster corrected). There were no statistically significant differences between the adult-onset blind and the congenitally blind groups in this contrast (Figure 2).

#### ROI analysis

In the LangOccip ROI, which included secondary visual areas, the sentences > non-words effect was larger in the congenitaly blind group than the sighted group (group-by-condition ANOVA CB vs S group-by-condition interaction F(1,38) = 8.28, p = 0.006), larger in the congenitally blind relative to the adult-onset blind group (group-by-condition ANOVA, AB vs. CB, group-by-condition interaction F(1,35) = 5.22, p = 0.028) and not significantly different between sighted and adult onset blind groups (group-by-condition ANOVA AB vs S group-by-condition interaction F(1,31) = 0.26, p = 0.616).

In V1, the response to sentences was larger in the congenitally blind relative to the adult-onset blind group (LangV1 group-by-condition ANOVA CB vs AB main effect of group F(1,35) = 6.17, p = 0.018 group-by-condition interaction F(1,35) = 3.43, p = 0.072), and the adult-onset blind showed a trending difference from the sighted group (group-by-condition ANOVA AB vs S group-by-condition interaction F(1,31) = 3.55, p = 0.068), suggesting an intermediate response profile.

In post-hoc within group comparisons, there was a significant difference between sentences and nonwords in the LangOccip ROI in the congenitally blind group (t(21) = 7.32, p < 0.001), the adult-onset blind group (t(14) = 4.54, p < 0.001) and in the sighted group (t(17) = 4.07, p < 0.001). In V1, there was a significant response to sentences > nonwords in the congenitally blind (t(21) = 5.01, p < 0.001) and adult-onset blind groups (t(14) = 4.29, p < 0.001), but not in the sighted group (t(17) = 1.25, p = 0.226).

In sum, the adult-onset blind group showed higher responses to sentences than non-words in both secondary visual areas and primary visual cortex, but this effect was smaller than what is observed in congenital blindness (Figure 3).

### 2.3 Sensitivity to syntactic complexity in visual cortex of congenitally blind but not adult-onset blind individuals (Experiment 2)

There was a significant movement-by group-interaction in both V1 and secondary visual cortex regions between the adult-onset blind and congenitally blind groups (movement-by-group ANOVA AB vs CB, movement-by-group interaction: LangV1: F(1,35) = 6.61, p = 0.015; LangOccip: F(1,35) = 6.25, p = 0.017). By contrast, there were no differences between the adult-onset blind and sighted groups with respect to the movement effect in any visual cortex regions (movement-by-group ANOVA, movement-by-group interaction LangOccip: F(1,31) = 1.18, p = 0.284; LangV1: F (1,31) = 0.43, p = 0.514) (Figure 3).

The same pattern held when we examined responses to syntactic movement in the region specifically responsive to sentences more than non-words in the adult-onset blind group (AB LangOccip). The adult-onset blind participants were no different from the sighted participants in their response to syntactic movement (movement-by-group ANOVA, movement-by-group interaction F(1,31) = 0.34, p = 0.559) in this region, and significantly differed from the congenitally blind (movement-by-group ANOVA, movement-by-group interaction F(1,35) = 13.95, p < 0.001).

There was a syntactic movement effect in the language-responsive secondary visual areas and V1 of congenitally blind adults (LangOccip: t(21) = 3.48, p = 0.002, LangV1 t(21) = 3.20, p = 0.004), and no effect of syntactic movement in the sighted participants (LangOccip: t(17) = 1.23, p = 0.234, LangV1 t(17) = −0.24, p = 0.813). The adult-onset blind participants patterned like the sighted group on this measure, i.e. there were no effects of syntactic movement in either LangOccip (t(14) = −0.47, p = 0.646) or in V1 (t(14) = −0.91, p = 0.378). Again, in the language-responsive region of the adult-onset blind group (AB LangOccip), there was a significant movement effect in the congenitally blind group (t(21) = 3.74, p = 0.001), but not in the adult-onset blind (t(14) = −1.78, p = 0.101) or the sighted group (t(17) = −1.18, p = 0.253).

To ensure that we were not missing a small effect in the adult-onset blind group, we repeated the analysis at smaller ROI sizes (top 10%, 5% and top 20 vertices). The adult-onset blind group failed to show a syntactic movement effect in any ROI, regardless of ROI size (all t’s < 1.0, all p’s > 0.1).

### 2.4. Relationship of blindness duration and age of blindness onset to visual cortex responses to language (Experiments 1 and 2)

In the adult onset-blind group, there was a tendency for responses to language in the visual cortex to increase with duration of blindness. Blindness duration and age of onset predicted the size of the sentences>non-words effect (Experiment 2) in the adult onset blind group in V1 (LangV1, duration: r = 0.63, t(13) = 2.94, p = 0.011, age of onset: r = −0.54, t(13) = −2.27, p = 0.040). When blindness duration and onset were both entered into a multiple regression, only the effect of blindness duration remained significant (LangV1, duration t(12) = 2.32, p = 0.037, blindness onset t(12) = −1.01, p = 0.327, adjusted r = 0.41). No other correlations were significant in any ROI, although all effects were in the same direction (p’s > 0.1) (Figure 4).

**Figure 4:**
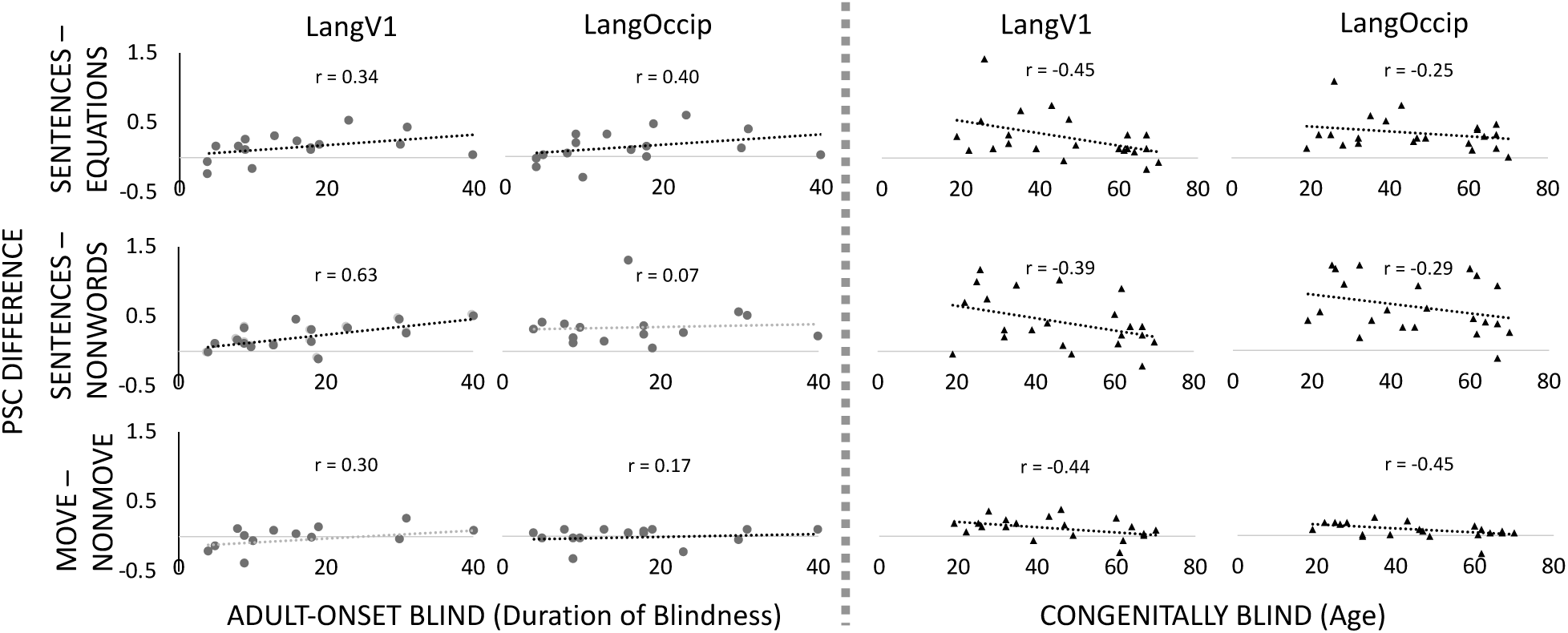
Effect of blindness duration (in Adult Onset Blind) and age (in Congenitally Blind) on visual cortex responses to language. PSC extracted from functionally defined individual LangV1 and CB LangOccip ROIs for the SENTENCE-NONWORD and MOVE-NONMOVE conditions in Experiment 2, and LangV1 and AB LangOccip ROIs for the SENTENCE-EQUATION conditions of Experiment 1.

By contrast, in the congenitally blind group, the response to language in visual cortex tended to decrease with blindness duration (i.e. age). This correlation was only significant in V1 for Experiment 1 (LangV1 r = −0.45, t(20) = −2.25, p = 0.036), and trending in V1 for Experiment 2 (LangV1 r = −0.39, t(20) = −1.88, p = 0.075), but was in the same direction in all comparisons (p’s > 0.1) (Figure 4).

The size of the MOVE-NONMOVE effect in both ROIs in the adult-onset blind group was not significantly predicted by duration of blindness (LangOccip: r = 0.17, t(13) = 0.63, p = 0.537; LangV1: r = 0.30, t(13) = 1.15, p = 0.272) or age of blindness onset (LangOccip: r = 0.36, t(13) = 1.40, p = 0.184; LangV1: r = 0.24, t(13) = 0.91, p = 0.379). In the congenitally blind group, the MOVE-NONMOVE effect showed a significant reduction in effect size with age in all visual cortex ROIs (LangOccip: r = −0.45, t(20) = −2.26, p = 0.035; LangV1: r = −0.44, t(20) = −2.17, p = 0.042).

### 2.5 Similar responses to language among sighted, congenitally blind and adult onset blind groups in inferior frontal cortex

The inferior frontal gyrus (IFG) showed a similar response profile in the adult-onset blind group relative to the sighted and congenitally blind groups (Figure 3). The IFG responded more to sentences than nonwords across groups (AB vs. CB main effect of condition F(1,35) = 198.91, p < 0.001, group-by-condition interaction F(1,35) = 0.02, p = 0.877; AB vs. S main effect of condition F(1,31) = 262.51, p < 0.001, group-by-condition interaction F(1,31) = 0.33, p = 0.567). There was also a larger response to the MOVE than NONMOVE sentences across groups (IFG AB vs. CB, main effect of condition F(1,35) = 12.18, p = 0.002, group-by-condition interaction F(1,35) = 0.61, p = 0.442; AB vs. S main effect of condition F(1,31) = 13.35, p < 0.001, group-by-condition interaction F(1,31) = 0.84, p = 0.365).

In post-hoc t-tests within the IFG, the sentences > nonwords effect was present in the congenitally blind (t(21) = 9.63, p < 0.001), adult-onset blind (t(14) = 10.98, p < 0.001) as well as the sighted group (t(17) = 11.94, p < 0.001). The syntactic movement effect was also present in all three groups (CB: t(21) = 3.67, p = 0.002; AB: t(14) = 1.79, p = 0.055; S: t(17) = 4.15, p < 0.001).

In whole-brain analyses, all three groups showed comparable fronto-temporal responses to sentences>math equations (Experiment 1) and sentences>nonwords (Exeriment 2) (Figure 2). However, fronto-temporal responses were left-lateralized in the sighted and adult-onset blind groups, but not in the congenitally blind group.

### 2.6 Reduced left lateralization of fronto-temporal reponses to language in congenitally but not adult-onset blind individuals

When we directly tested the laterality of fronto-temporal language responses, we observed reduced left-lateralization relative to the sighted in the congenitally but not adult-onset blind group (Figure 5). The response to spoken language in the fronto-temporal language network of the adult-onset blind group was as left-lateralized as in the sighted, both in the sentences > math contrast (t(13) = 0.14, p = 0.889) and the sentences > non-words contrast (t(13) = −0.19, p = 0.846). The adult-onset blind group was significantly more left lateralized than the congenitally blind (sentences > mathematics t(13) = −3.33, p = 0.002; sentences > nonwords t(13) = −2.56, p = 0.015).

**Figure 5:**
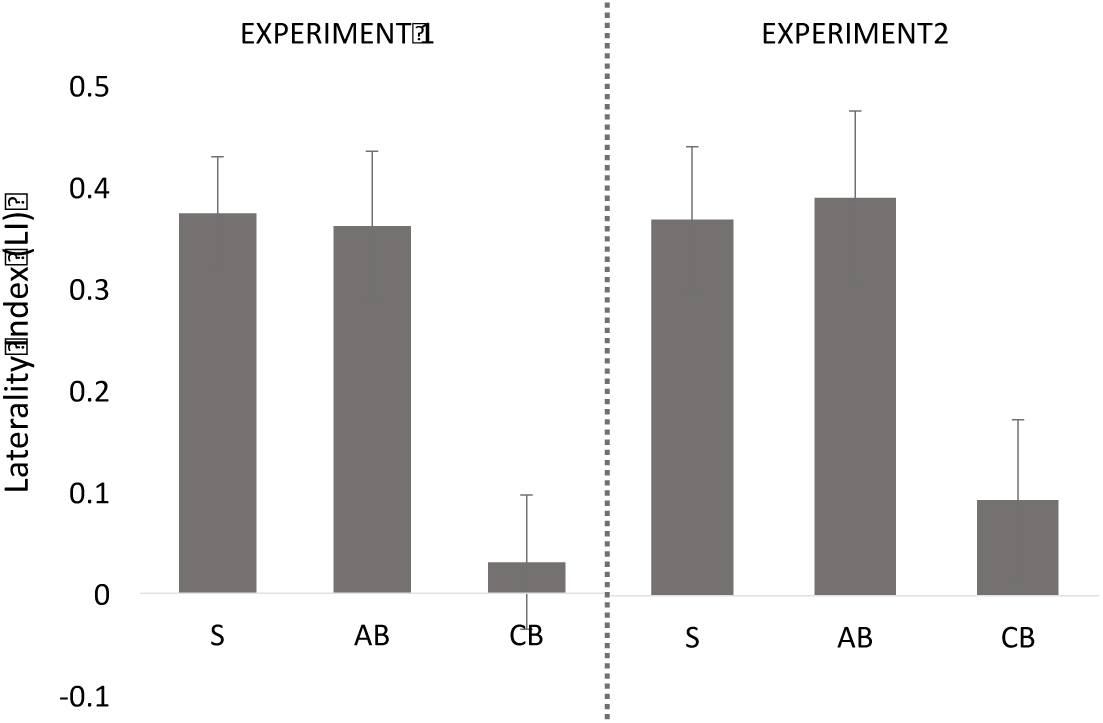
Mean laterality index for sighted (S), adult-onset blind (AB) and congenitally blind (CB) participants in response to the sentences > mathematics (Experiment 1) and sentences > nonwords (Experiment 2) condition in the whole brain excluding the occipital cortex. Error bars represent SEM.

This reduction of left lateralization was equivalent when we separated the CB group based on ROP (59%) and non-ROP as cause of blindness. There was no correlation between Braille reading scores and the laterality index in either group, in Experiment 1 or 2 (all p’s > 0.1). There was also no correlation in either group or experiment between laterality index and duration of blindness (all p’s > 0.1).

## DISCUSSION

### A sensitive period in the neural substrates of language in blindness

Previous studies identified two ways in which the neural basis of language processing is modified in blind individuals. First, parts of the “visual” cortex are incorporated into the language network and become sensitive to the grammatical structure of spoken sentences (Bedny et al., 2011; Lane et al., 2015; Röder et al., 2002). Second, fronto-temporal language areas are less left-lateralized in congenitally blind than sighted individuals (Lane et al., 2017). Here, we report that both of these phenomena follow a sensitive period in development and are either absent, or present to a much lesser extent in individuals who lose their vision as adults.

Adult-onset blind individuals show left-lateralization of fronto-temporal language areas that is indistinguishable from sighted adults, and greater than congenitally blind participants. “Visual” cortices of adult-onset blind individuals differentiate less between sentences and equations and sentences and lists of non-words than those of congenitally blind individuals, although some difference between these conditions is observed even in individuals who became blind later in life. We observed sensitivity to syntactic movement in “visual” cortex in congenitally blind, but not adult-onset blind adults. These results are consistent with the idea that blindness has different effects on the neural basis of language during and after a developmental sensitive period, and therefore support the developmental specialization hypothesis.

The absence of a syntactic movement effect in the “visual” cortex in adult-onset blindness is particularly intriguing, and consistent with claims that sensitivity to aspects of syntax depends on cortical flexiblity that is inherent to sensitive periods (Friederici, 2017; Lenneberg, 1967; Neville et al., 1992; Ruben, 1999). Studies of delayed first language exposure, as well as second language learning provide support for this hypothesis (Cormier, Schembri, Vinson, & Orfanidou, 2012; Mayberry, 2007; Mayberry & Eichen, 1991; Mayberry & Lock, 2003). For example, several studies suggest that hearing impaired children fitted with cochlear implants later in life show significant deficits in morpho-syntactic production and comprehension tasks, while still being comparable in performance on vocabulary and lexical semantic tasks to age matched hearing children (Friedmann & Rusou, 2015; Geren & Snedeker, 2009; Lopez-Higes, Gallego, Martin-Aragoneses, & Melle, 2015). Indeed, there is some evidence that syntactic movement per se is affected by delays in language access (Friedmann, 2005; Friedmann & Rusou, 2015). In a neuroimaging study of grammaticality and phonemic judgements, age of acquisition also affected the neurobiology of sign language, with later learners of ASL showing lower BOLD responses to syntactic tasks in left inferior frontal regions (Mayberry et al., 2011). Analogously, second language speakers who aquire a language as adults show a reduced ability to acquire aspects of grammatical structure in their second language, while having less trouble with vocabulary acquisition and basic word order (Johnson & Newport, 1989; Newport, 1990; Patkowski, 1980; Weber-Fox & Neville, 1996). A recent large-scale study with over half a million participants has confirmed age-of-acqusition effects on sentence-level syntax in second language learners, although it also suggests that in this particular case, the cortical learning rate may not fall off until the late teens (Hartshorne, Tenenbaum, & Pinker, 2018). There is also evidence of different neural signatures for syntax in first and second languages speakers matched for proficiency (Neville et al., 1998; Pakulak & Neville, 2011; Weber-Fox & Neville, 2001).

The present results provide complementary evidence for the hypothesis that acquisition of aspects of grammar depends on the special cortical flexibility afforded by sensitive periods. One interpretation of previous sensitive period effects in grammar is that they arise uniquely from the maturational timetable characteristic to fronto-temporal language areas (Lenneberg, 1967; Newport & York, 2003). An alternative, non-mutually exclusive possiblity suggested by the present findings is that certain aspects of syntax acquisition depend on critical period plasticity more generally, even outside the fronto-temporal network.

### Visual cortex of adult-onset blind individuls responds to spoken sentences

Although we observed different responses to spoken sentences in visual cortices of congenitally and adult-onset blind individuals, we found that in adult-onset blind individuals, both V1 and secondary “visual” areas respond to spoken sentences more than to lists of non-words and equations. In some cases, small effects were observed even in the secondary visual areas of the blindfolded sighted group. Furthermore, in the adult-onset blind group, there was a tendency for responses to spoken language to increase over the course of many years, with AB individuals who are blind for 30 years showing larger responses to sentences than those who are blind for 20. Thus, sensitive period effects coexist with continued plasticity throughout life. One interpretation is that the “visual” cortex retains the capacity for functional reorganization into adulthood, but in adulthood, the rate of learning is much slower (Merabet & Pascual-Leone, 2010). Within this framework, lack of responses to syntactic movement in the visual cortex of adult-onset blind individuals might reflect the fact that the human lifespan is insufficiently long to acquire such sensitivity, given the slower learning rate in adult cortex.

The presence of responses to language in the visual cortex in individuals who become blind as adults is consistent with the observation of increased resting-state connectivity between Broca’s area and the “visual” cortex in this population (Sabbah et al., 2016). Analogously, a recent study found increased resting-state connectivity between parts of the “visual” cortex that are responsive to number, and fronto-parietal number networks, even in adult-onset blind individuals (Kanjlia et al., 2018). This latter study also showed that resting-state increases are significantly smaller in the adult-onset as opposed to the congenitally blind population. Furthermore, like in the current study, sensitivity to task-based cognitive manipulations was reduced or absent in adult-onset blindness - in the case of this prior study, responses to the difficulty of math equations were present only in people blind from birth. One possibility is that acquisition of sensitivity to fine-grained cognitive distinctions (e.g. to syntax and equation difficulty) depends on critical period plasticity in local cortical circuits (Hensch, 2005).

Previous studies also suggest that the behavioral relevance of “visual” cortex activity in adult onset and congenital blindness is different. In congenital blindness, transcranial magnetic stimulation (TMS) to the occipital pole induces semantic errors in a verb generation task (Amedi, Floel, Knecht, Zohary, & Cohen, 2004). Congenitally blind adults also show superior performance on some linguistic tasks that recruit the “visual” cortex e.g. verbal memory (Amedi, Raz, Pianka, Malach, & Zohary, 2003; Occelli, Lacey, Stephens, Merabet, & Sathian, 2017; Pasqualotto, Lam, & Proulx, 2013). By contrast, there is at present no evidence for functional relevance of visual cortices to language (or any other cognitive function) in adult-onset blindness. Indeed, one study found that TMS to the occipital pole impairs Braille reading in congenitally blind, but not adult-onset blind individuals (Cohen et al., 1999). It thus remains possible that activity in visual cortices of adult-onset blind individuals is epiphenomenal with respect to behavior. This could be because the degree of involvement of the “visual” cortex is so small in the adult-onset blind population as to be task irrelevant. It also remains possible, however, that cross-modal “visual” cortex activity in adult onset blind individuals has a different cognitive role from that of congenitally blind individuals, and this role has not yet been tested.

Even if responses to language in the visual system of adult-onset blind individuals are not behaviorally relevant, they support the hypothesis that communication between visual and language systems exists even in individuals whose brain developed with vision (Tomasello, Garagnani, Wennekers, & Pulvermüller, 2019). As noted above, we observed small but reliable effects in the secondary but not primary visual areas of sighted blindfolded individuals as well, i.e. the sighted group showed significantly higher responses to sentences than non-words in high-level visual areas. Previous studies have also observed some seemingly paradoxical sensitivity to language in the “visual” system of blindfolded sighted adults. One study of noun and verb processing found elevated responses to verbs relative to nouns in lateral occipital cortices of blindfolded sighted individuals (Elli et al., 2019). Notably, overall responses to nouns and verbs were below rest in these visual areas, and unlike in temporo-parietal networks, visual regions of the sighted were not sensitive to semantic distinctions among nouns or verbs in MVPA analysis (Elli et al., 2019). These results suggest that communication exists between the language and the “visual” networks even in sighted adults. One interpretation is that blindness from birth modifies the way this information is used by “visual” cortex.

How might language information get to the visual system? In humans, the inferior fronto-occipital fasciculus (IFOF) contains a set of fibers passing from the occipital lobe to the inferior frontal cortex, the inferior longitudinal fasciculus (ILF) connects the occipital to the anterior and medial temporal lobe, and the vertical occipital fasciculus (VOF) of Wernicke connects the ventral occipitotemporal cortex to the lateral occipito-parietal junction (Ashtari, 2012; Forkel et al., 2014; Yeatman, Rauschecker, & Wandell, 2013). In the sighted, these connections may enable task specific interactions between vision and language, such interactions might occur during tasks such as describing a visual scene, identifying objects based on verbal labels, or retrieving abstract linguistic representations from visual symbols i.e. reading (Jackendoff, 1987; Landau & Jackendoff, 2013).

The visual word form area (VWFA) is one of the key cortical nodes that connects visual and language systems (Bouhali et al., 2014; Yeatman et al., 2014). In sighted individuals, this region develops selectivity for written language, putatively because its function is to connect visual symbols with linguistic content (Dehaene & Dehaene-Lambertz, 2016; Saygin et al., 2016). In blindness, the VWFA responses to written (Braille) and spoken language (Büchel, Price, & Friston, 1998; Burton et al., 2002; Cohen et al., 1997; Reich, Szwed, Cohen, & Amedi, 2011; Röder et al., 2002; Sadato, 2005). Recent evidence further suggests that the VWFA develops sensitivity to grammar in people blind from birth (Kim et al., 2017). Object-responsive lateral occipital areas are another potential point of contact between visual and linguistic systems because of their anatomical proximity to the language system (Connolly et al., 2012; Kanwisher, 2010; Osher et al., 2016). In blind participants, the lateral occipital cortex shows responses to high-level linguistic information (Bedny et al., 2011; Kim et al., 2017; Lane et al., 2015; Röder et al., 2002). Ventral and lateral occipito-temporal areas may serve as entry points for linguistic information into the visual system. In this regard, one might view blindness-related plasticity as a special instance of cortical map expansion, where in this case language expands into territory typically occupied by visual functions (Jones, 2002). Notably, responses to language in “visual” cortex of blind individuals are also observed in retinotopic regions, including V1 (Bedny et al., 2011; Lane et al., 2015; Watkins et al., 2012). Whether linguistic information first enters “visual” networks via secondary visual areas and feedback to V1, or whether there is a separate route to primary visual cortex remains to be determined.

### Summary and Conclusions

The present results reveal a sensitive period for the reorganization of language networks in blindness. Only in congenitally blind individuals do visual cortices respond to syntactic movement, and visual cortex responses to spoken sentences are much larger in congenitally than adult-onset blind individuals. These results are consistent with the idea that in the absence of dominating visual input from the lateral geniculate nucleus, parts of the visual system are incorporated into the language network during language acquisition. The plasticity observed in congenital blindness supports the idea that the neural basis of language, while evolutionarily constrained, nevertheless emerges through a dynamic process that includes competition for the same cortical territory by multiple cognitive functions (Bates, 1993; Johnson, Halit, Grice, & Karmiloff-Smith, 2002; Karmiloff-Smith, 1998). The presence of some high-level language responses even in the visual system of adult-onset blind and blindfolded sighted people suggests that the plasticity observed in congenital blindness is made possible by existing channels of communication between the visual and language systems.

The current results add to prior evidence of different cognitive sensitivity in the visual cortices of congenitally and adult-onset blind individuals (eg: Bedny, Konkle, Pelphrey, Saxe, & Pascual-Leone, 2010; Bedny, Pascual-Leone, Dravida, & Saxe, 2012; Büchel, Price, Frackowiak, & Friston, 1998; Burton, Diamond, & McDermott, 2006; Burton et al., 2002; Cohen et al., 1999; Kanjlia, Pant, & Bedny, 2018). Previous studies have found differences in visual cortex recruitment between late onset and congenital blindness in auditory spatial/pitch processing (Collignon et al., 2013), auditory motion perception (Bedny et al., 2010), monaural and binaural auditory localization tasks (Voss, Gougoux, Zatorre, Lassonde, & Lepore, 2008), in tactile discrimination (Cohen et al., 1999) and numerical cognition (Kanjlia et al., 2018). Together with the present results, these studies support the hypothesis that human cortex has a different capacity for cognitive specialization during childhood, as opposed to in adulthood.

## Acknowledgements

We thank the blind and sighted participants for taking part in this research. We are also grateful to Connor Lane and Dr. Akira Omaki for assistance with the design of the stimuli and task in Experiment 2. This study was supported by grant R01EY027352-02 from the National Institute of Health to Dr. Marina Bedny.

## Supplementary Material

### S1: Peak table

*Language responsive regions in Sighted, Adult-Onset Blind and Congenitally Blind individuals. X,Y and Z values are in MNI space. Peak t values are for the local minima, and all p values are cluster corrected. All peaks reported are at least 20 mm apart.*

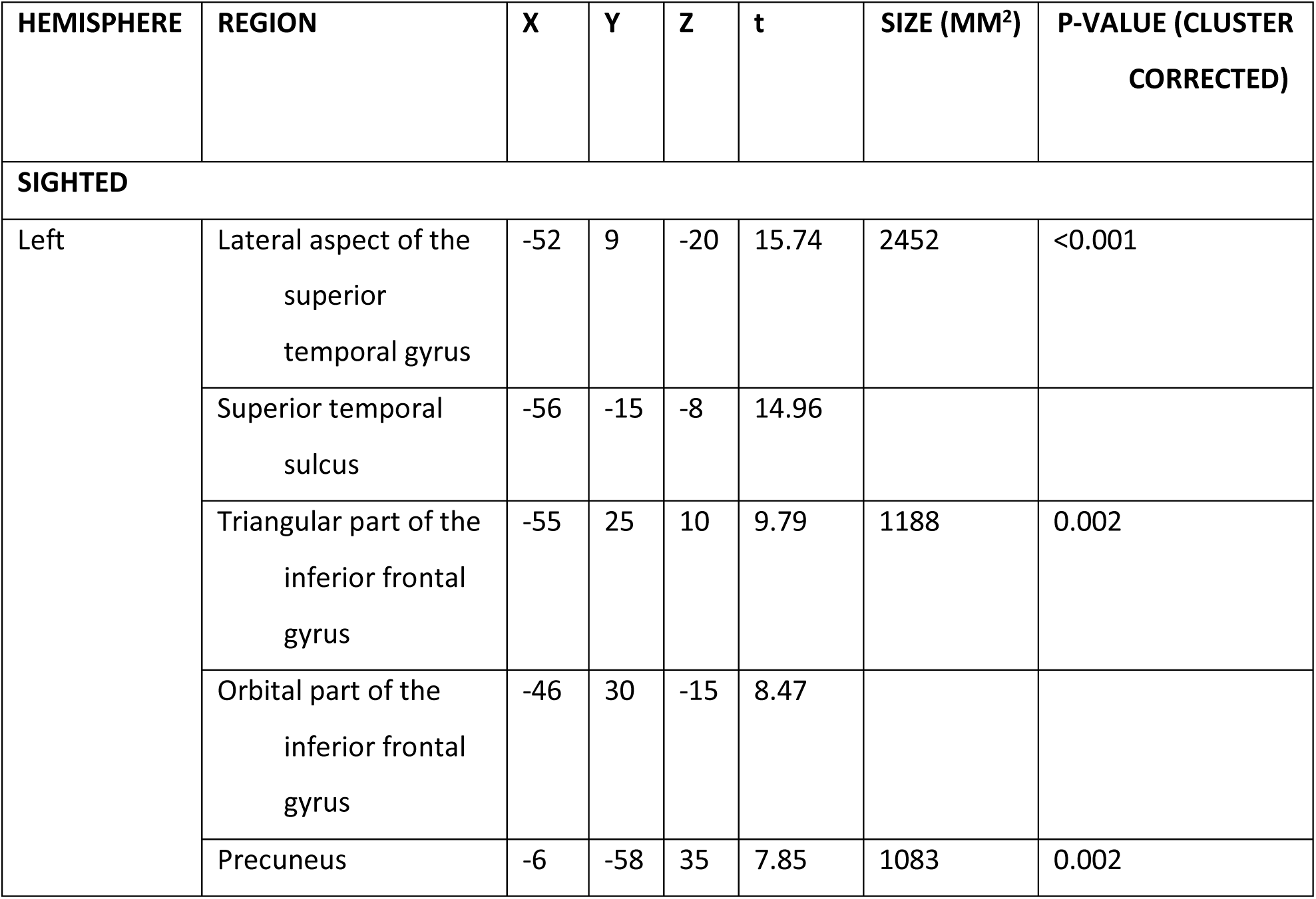
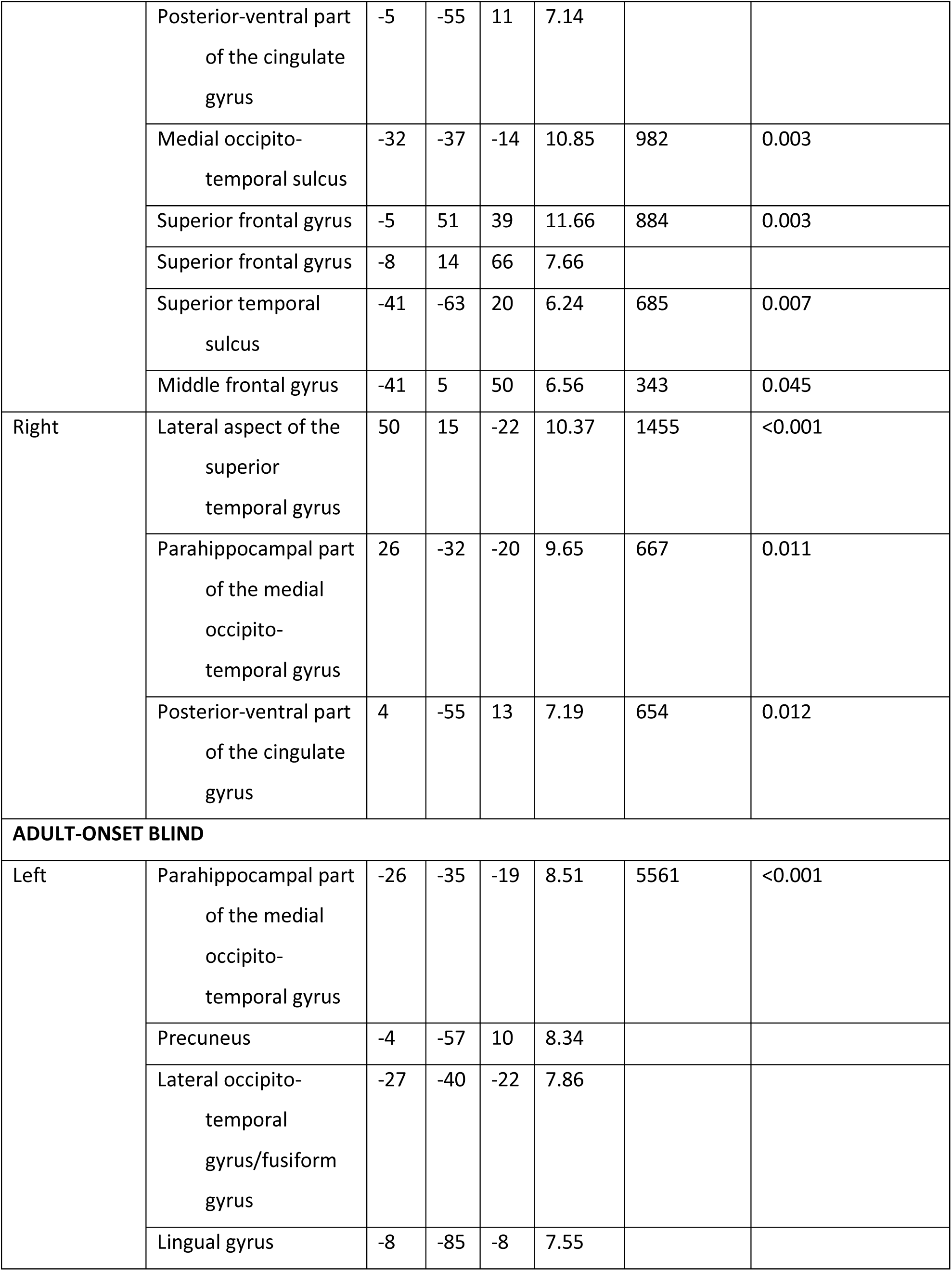
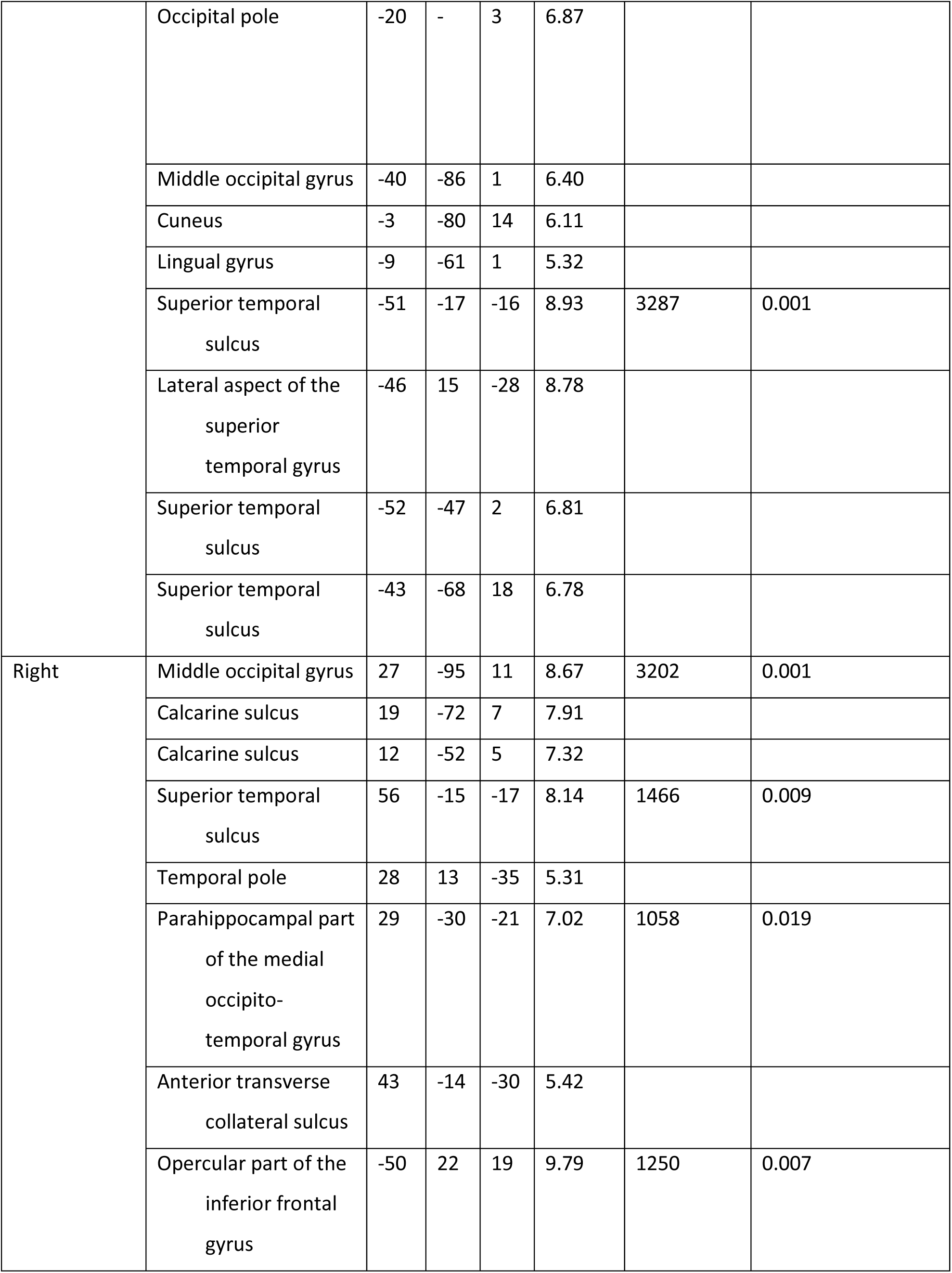
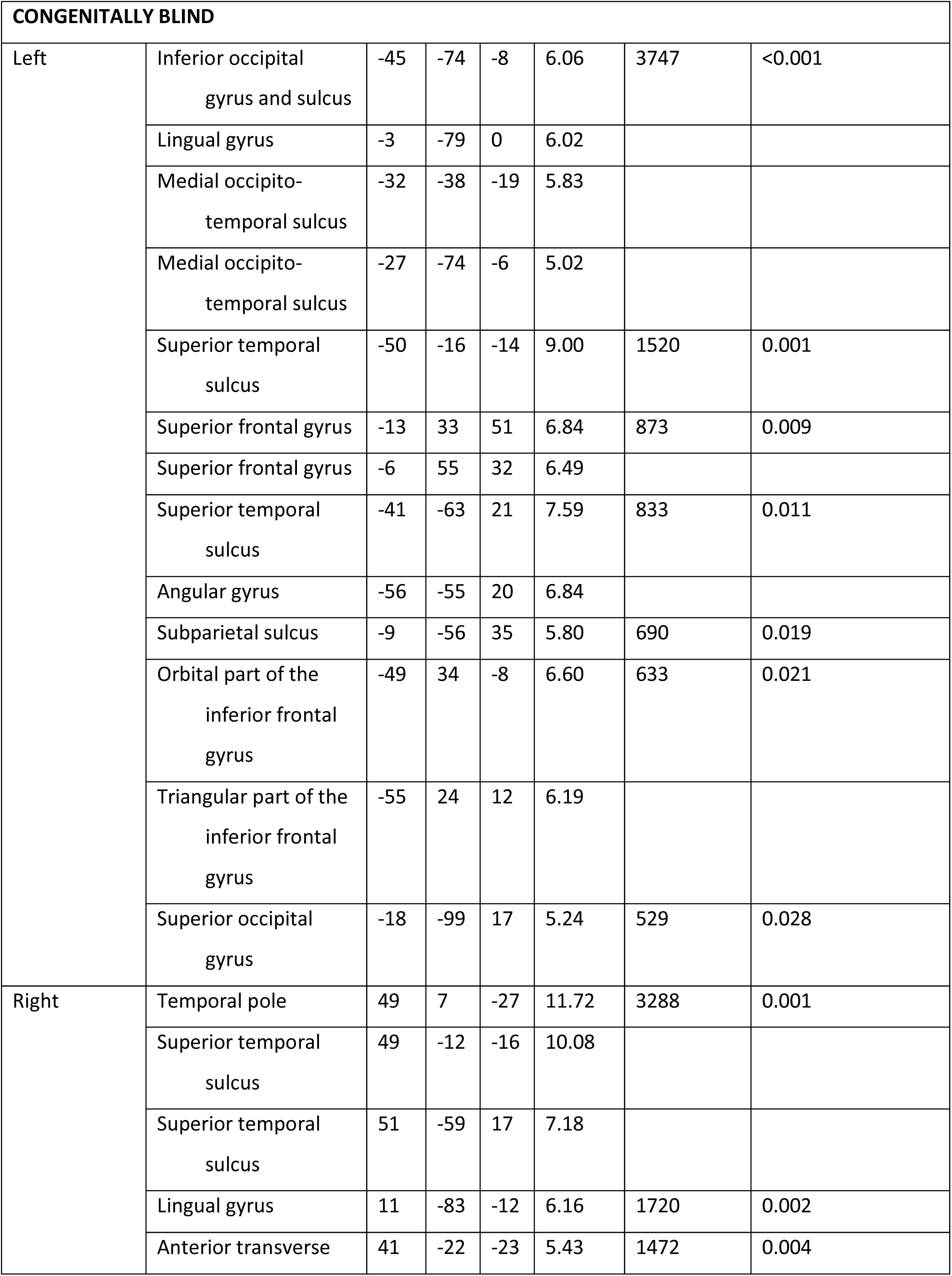
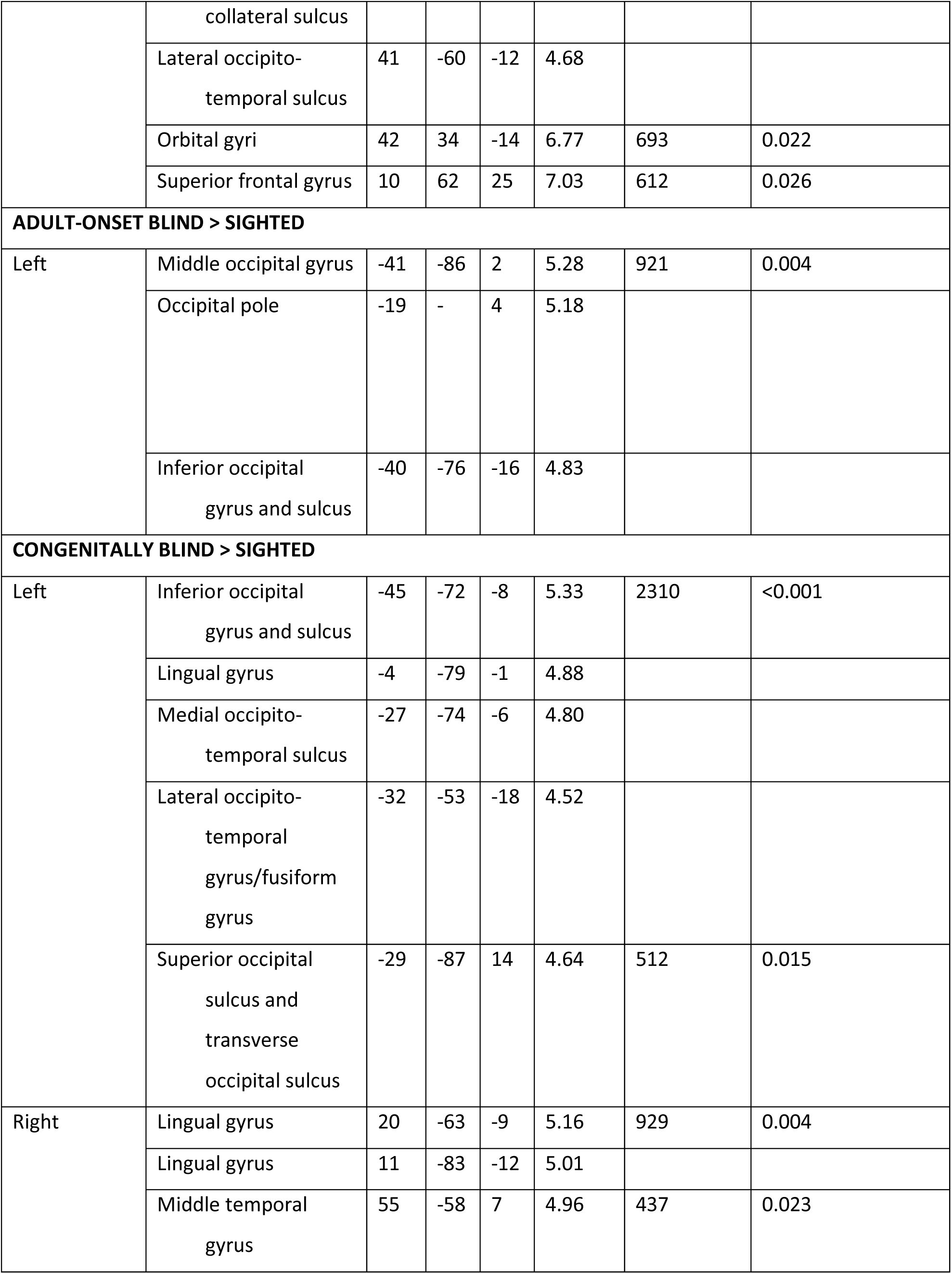

| HEMISPHERE | REGION | X | Y | Z | t | SIZE (MM<sup>2</sup>) | P-VALUE (CLUSTER CORRECTED) |
| --- | --- | --- | --- | --- | --- | --- | --- |
| **SIGHTED** |  |  |  |  |  |  |  |
| Left | Lateral aspect of the superior temporal gyrus | -52 | 9 | -20 | 15.74 | 2452 | <0.001 |
|  | Superior temporal sulcus | -56 | -15 | -8 | 14.96 |  |  |
|  | Triangular part of the inferior frontal gyrus | -55 | 25 | 10 | 9.79 | 1188 | 0.002 |
|  | Orbital part of the inferior frontal gyrus | -46 | 30 | -15 | 8.47 |  |  |
|  | Precuneus | -6 | -58 | 35 | 7.85 | 1083 | 0.002 |
|  | Posterior-ventral part of the cingulate gyrus | -5 | -55 | 11 | 7.14 |  |  |
|  | Medial occipito-temporal sulcus | -32 | -37 | -14 | 10.85 | 982 | 0.003 |
|  | Superior frontal gyrus | -5 | 51 | 39 | 11.66 | 884 | 0.003 |
|  | Superior frontal gyrus | -8 | 14 | 66 | 7.66 |  |  |
|  | Superior temporal sulcus | -41 | -63 | 20 | 6.24 | 685 | 0.007 |
|  | Middle frontal gyrus | -41 | 5 | 50 | 6.56 | 343 | 0.045 |
| Right | Lateral aspect of the superior temporal gyrus | 50 | 15 | -22 | 10.37 | 1455 | <0.001 |
|  | Parahippocampal part of the medial occipito-temporal gyrus | 26 | -32 | -20 | 9.65 | 667 | 0.011 |
|  | Posterior-ventral part of the cingulate gyrus | 4 | -55 | 13 | 7.19 | 654 | 0.012 |
| **ADULT-ONSET BLIND** |  |  |  |  |  |  |  |
| Left | Parahippocampal part of the medial occipito-temporal gyrus | -26 | -35 | -19 | 8.51 | 5561 | <0.001 |
|  | Precuneus | -4 | -57 | 10 | 8.34 |  |  |
|  | Lateral occipito-temporal gyrus/fusiform gyrus | -27 | -40 | -22 | 7.86 |  |  |
|  | Lingual gyrus | -8 | -85 | -8 | 7.55 |  |  |
|  | Occipital pole | -20 | - | 3 | 6.87 |  |  |
|  | Middle occipital gyrus | -40 | -86 | 1 | 6.40 |  |  |
|  | Cuneus | -3 | -80 | 14 | 6.11 |  |  |
|  | Lingual gyrus | -9 | -61 | 1 | 5.32 |  |  |
|  | Superior temporal sulcus | -51 | -17 | -16 | 8.93 | 3287 | 0.001 |
|  | Lateral aspect of the superior temporal gyrus | -46 | 15 | -28 | 8.78 |  |  |
|  | Superior temporal sulcus | -52 | -47 | 2 | 6.81 |  |  |
|  | Superior temporal sulcus | -43 | -68 | 18 | 6.78 |  |  |
| Right | Middle occipital gyrus | 27 | -95 | 11 | 8.67 | 3202 | 0.001 |
|  | Calcarine sulcus | 19 | -72 | 7 | 7.91 |  |  |
|  | Calcarine sulcus | 12 | -52 | 5 | 7.32 |  |  |
|  | Superior temporal sulcus | 56 | -15 | -17 | 8.14 | 1466 | 0.009 |
|  | Temporal pole | 28 | 13 | -35 | 5.31 |  |  |
|  | Parahippocampal part of the medial occipito-temporal gyrus | 29 | -30 | -21 | 7.02 | 1058 | 0.019 |
|  | Anterior transverse collateral sulcus | 43 | -14 | -30 | 5.42 |  |  |
|  | Opercular part of the inferior frontal gyrus | -50 | 22 | 19 | 9.79 | 1250 | 0.007 |

| CONGENITALLY BLIND |  |  |  |  |  |  |  |
| --- | --- | --- | --- | --- | --- | --- | --- |
| Left | Inferior occipital gyrus and sulcus | -45 | -74 | -8 | 6.06 | 3747 | <0.001 |
|  | Lingual gyrus | -3 | -79 | 0 | 6.02 |  |  |
|  | Medial occipito-temporal sulcus | -32 | -38 | -19 | 5.83 |  |  |
|  | Medial occipito-temporal sulcus | -27 | -74 | -6 | 5.02 |  |  |
|  | Superior temporal sulcus | -50 | -16 | -14 | 9.00 | 1520 | 0.001 |
|  | Superior frontal gyrus | -13 | 33 | 51 | 6.84 | 873 | 0.009 |
|  | Superior frontal gyrus | -6 | 55 | 32 | 6.49 |  |  |
|  | Superior temporal sulcus | -41 | -63 | 21 | 7.59 | 833 | 0.011 |
|  | Angular gyrus | -56 | -55 | 20 | 6.84 |  |  |
|  | Subparietal sulcus | -9 | -56 | 35 | 5.80 | 690 | 0.019 |
|  | Orbital part of the inferior frontal gyrus | -49 | 34 | -8 | 6.60 | 633 | 0.021 |
|  | Triangular part of the inferior frontal gyrus | -55 | 24 | 12 | 6.19 |  |  |
|  | Superior occipital gyrus | -18 | -99 | 17 | 5.24 | 529 | 0.028 |
| Right | Temporal pole | 49 | 7 | -27 | 11.72 | 3288 | 0.001 |
|  | Superior temporal sulcus | 49 | -12 | -16 | 10.08 |  |  |
|  | Superior temporal sulcus | 51 | -59 | 17 | 7.18 |  |  |
|  | Lingual gyrus | 11 | -83 | -12 | 6.16 | 1720 | 0.002 |
|  | Anterior transverse collateral sulcus | 41 | -22 | -23 | 5.43 | 1472 | 0.004 |
|  | Lateral occipito-temporal sulcus | 41 | -60 | -12 | 4.68 |  |  |
|  | Orbital gyri | 42 | 34 | -14 | 6.77 | 693 | 0.022 |
|  | Superior frontal gyrus | 10 | 62 | 25 | 7.03 | 612 | 0.026 |
| **ADULT-ONSET BLIND > SIGHTED** |  |  |  |  |  |  |  |
| Left | Middle occipital gyrus | -41 | -86 | 2 | 5.28 | 921 | 0.004 |
|  | Occipital pole | -19 | - | 4 | 5.18 |  |  |
|  | Inferior occipital gyrus and sulcus | -40 | -76 | -16 | 4.83 |  |  |
| **CONGENITALLY BLIND > SIGHTED** |  |  |  |  |  |  |  |
| Left | Inferior occipital gyrus and sulcus | -45 | -72 | -8 | 5.33 | 2310 | <0.001 |
|  | Lingual gyrus | -4 | -79 | -1 | 4.88 |  |  |
|  | Medial occipito-temporal sulcus | -27 | -74 | -6 | 4.80 |  |  |
|  | Lateral occipito-temporal gyrus/fusiform gyrus | -32 | -53 | -18 | 4.52 |  |  |
|  | Superior occipital sulcus and transverse occipital sulcus | -29 | -87 | 14 | 4.64 | 512 | 0.015 |
| Right | Lingual gyrus | 20 | -63 | -9 | 5.16 | 929 | 0.004 |
|  | Lingual gyrus | 11 | -83 | -12 | 5.01 |  |  |
|  | Middle temporal gyrus | 55 | -58 | 7 | 4.96 | 437 | 0.023 |

